# ImmunoStruct: a multimodal neural network framework for immunogenicity prediction from peptide-MHC sequence, structure, and biochemical properties

**DOI:** 10.1101/2024.11.01.621580

**Authors:** Kevin Bijan Givechian, João Felipe Rocha, Edward Yang, Chen Liu, Kerrie Greene, Rex Ying, Etienne Caron, Akiko Iwasaki, Smita Krishnaswamy

**Author notes:** These authors contributed equally to this work (João Felipe Rocha, Edward Yang, Chen Liu).

## Abstract

Epitope-based vaccines are promising therapeutic modalities for infectious diseases and cancer, but identifying immunogenic epitopes is challenging. The vast majority of prediction methods only use amino acid sequence information, and do not incorporate wide-scale structure data and biochemical properties across each peptide-MHC. We present ImmunoStruct, a deep-learning model that integrates sequence, structural, and biochemical information to predict multi-allele class-I peptide-MHC immunogenicity. By leveraging a multimodal dataset of *∼*27,000 peptide-MHCs, we demonstrate that ImmunoStruct improves immunogenicity prediction performance and interpretability beyond existing methods, across infectious disease epitopes and cancer neoepitopes. We further show strong alignment with *in vitro* assay results for a set of SARS-CoV-2 epitopes, as well as strong performance in peptide-MHC-based cancer patient survival prediction. Overall, this work also presents a new architecture that incorporates equivariant graph processing and multimodal data integration for the long standing task in immunotherapy.

## Introduction

Epitope-based vaccines are a type of targeted immunotherapy that use short peptides to prime the immune system to attack infected or cancerous cells in the body. A peptide is considered to contain an immunogenic epitope if it can successfully invoke an immune response in the host, typically by activating T cells that target the same specific epitope. Epitope-based vaccines have shown promise in the clinic across cancer types such as melanoma, bladder carcinoma, glioblastoma, lung adenocarcinoma, and pancreatic cancer [1– 3]. However, vaccine delivery constructs, like lipid nanoparticles for mRNA vaccines, typically only hold up to 20-30 candidate peptides. Additionally, only a subset of patients respond to a limited fraction of these epitopes, and validating epitope immunogenicity remains resource-intensive [2–4]. While patients with observed CD8+ T-cell-mediated responses can exhibit encouraging clinical outcomes, improved tools are still needed for prioritizing and ranking epitope immunogenicity to enhance candidate selection. For instance, in previous analyses only 2-6% of predicted mutation-derived peptides were shown to actually result in true patient derived T-cell immunogenicity [4–7]. Additionally, recent cancer neoantigen vaccine trials have predominantly only used transcript expression and affinity-based methods for filtering and identifying patient-specific epitopes, resulting in a fraction of true positive predicitons [8]. Therefore, given the rarity of immunogenic epitopes and the paucity of reliable prediction methods, an important challenge remains in developing methods to distinguish immunogenic peptides from those that are not in order to enhance epitope-based vaccine design.

The surveilance of the human immune system relies on presentation of antigen peptides on top of major histocompatibility (MHC) proteins, which occurs via two distinct peptide presentation pathways – Class-I and Class-II. MHC Class-I proteins are found on most cells. These proteins present peptides to cytotoxic CD8+ T-cells, while MHC Class-II proteins are found on antigen-presenting cells and interact with helper-T-cells [9, 10]. A key goal in epitope-based vaccine design is to induce more robust CD8+ T-cell responses towards pathogenic cells, but pinpointing Class-I immunogenicity determinants is complex as it involves multiple cellular mechanisms. These include antigen processing, peptide-MHC binding, peptide-MHC presentation, peptide-MHC-TCR recognition, and CD8+ T-cell activation [9]. The same peptide can also show varying CD8+ T-cell activation based on its paired MHC allele, which vary significantly across the human population [10]. These variable peptide-MHC responses exist partly because structural differences in the peptide-MHC also influence binding cooperativity [11, 12]. Despite the development of numerous peptide-MHC deep-learning models like NetMHCpan-4.1 [13], MHCnuggets [14], MHCflurry-2.0 [15], and BigMHC [16], most still rely mainly on peptide and MHC sequence data without incorporating multi-allelic structural information and biochemical information [7], which are proven to be critical for peptide-MHC-TCR synapse recognition and immunogenicity [17].

To address these challenges, we present ImmunoStruct, a deep-learning model that integrates peptide-MHC sequence, structure, and biochemical properties to predict class-I peptide-MHC immunogenicity. To the best of our knowledge, this is the first method to leverage these combined modalities for immunogenicity prediction across both infectious disease and cancer.

ImmunoStruct is a multimodal neural network architecture, aiming to effectively integrate sequence information, structural peptide-MHC information, and biochemical property information across all samples to improve immunogenicity prediction. Immunostruct consists of a variational autoencoder (VAE) module that encodes a compressed representation of a peptide-MHC sequence, a graph transformer module that processes the graph constructed from both sequence and structure, and an MLP layer that encodes biochemical information. Further, Immunostruct features a multi-modal attention mechanism, that enables an innovative layer of structural interpretability over its predictions. In addition, given the availability of both wildtype and mutant peptide sequences in the cancer neoantigen context, ImmunoStruct uses a novel contrastive loss approach that leverages both the wildtype peptide-MHC data to enforce wildtype/mutant pairs to be further away in latent space if the mutant variant is immunogenic and vice versa. This helps the model distinguish whether a given mutation in a wildtype sequence would yield an immunogenic mutant peptide.

To train immunostruct, we custom-curated a multi-allelic dataset of peptide-MHC sequences using the IEDB [18] and CEDAR [19] databases – IEDB data were used for infectious diseases and CEDAR was used for cancer neoeptiopes. We then generated the peptide-MHC using our high-throughput AlphaFold2 pipeline [20] where each peptide-MHC pair was jointly-folded. Finally, biochemical properties were computed using both sequences and structures, creating our final multi-modal dataset, which in total contained the three distinct modalities.

ImmunoStruct was evaluated on held-out test data from both infectious disease and cancer from IEDB and CEDAR, respectively, showing consistently improved performance over current state-of-the-art models [13–16, 21, 22]. Moreover, given the importance of experimental validation especially for deep learning approaches, we generated an *in vitro* assay using ELISpot IFNg for a set of 19 peptide-MHCs. These peptides comprised various components of SARS-CoV-2 proteins with confirmed CD8+ T-cell immunogenicities. We also validated ImmunoStruct in a cohort of cancer patient survival comprising a variety of different tumor types and immunotherapeutic treatments. Together, with these experimental validation steps, ImmunoStruct also demonstrated clinical translation potential.

Overall, the key contributions of ImmunoStruct include the following:

- An multimodal neural network architecture capable of synergistically integrating three distinct data modalities for the immunogenicity prediction task.
- Curation of a multi-modal dataset enabling the use of structural, sequence, and biochemical property information to improve prediction of peptide-MHC immunogenicity.
- Interpretability of peptide-MHC predictions derived from attention-guided biostructural insights for immunogenic peptide-MHC samples.
- A new contrastive loss approach for the classification of cancer neoepitopes that forces wildtype and immunogenic mutant peptides to be embedded further away in latent space.

## Results

### The ImmunoStruct model

ImmunoStruct integrates protein sequence, structure, and biochemical information for peptide-MHC immunogenicity prediction (Figure 1a). The model takes in three input modalities: 1) peptide-MHC sequence, 2) peptide-MHC 3D structure given by AlphaFold2 (Figure 1b), and 3) biochemical properties (Figure 1c). These three modalities are modeled using a varational autoencoder (VAE), a graph transformer network, and a multilayer perceptron (MLP), which respectively encode the sequence, the structure/sequence combination, and the biochemical properties. The multimodal embedding is then passed to a downstream

**Figure 1:**
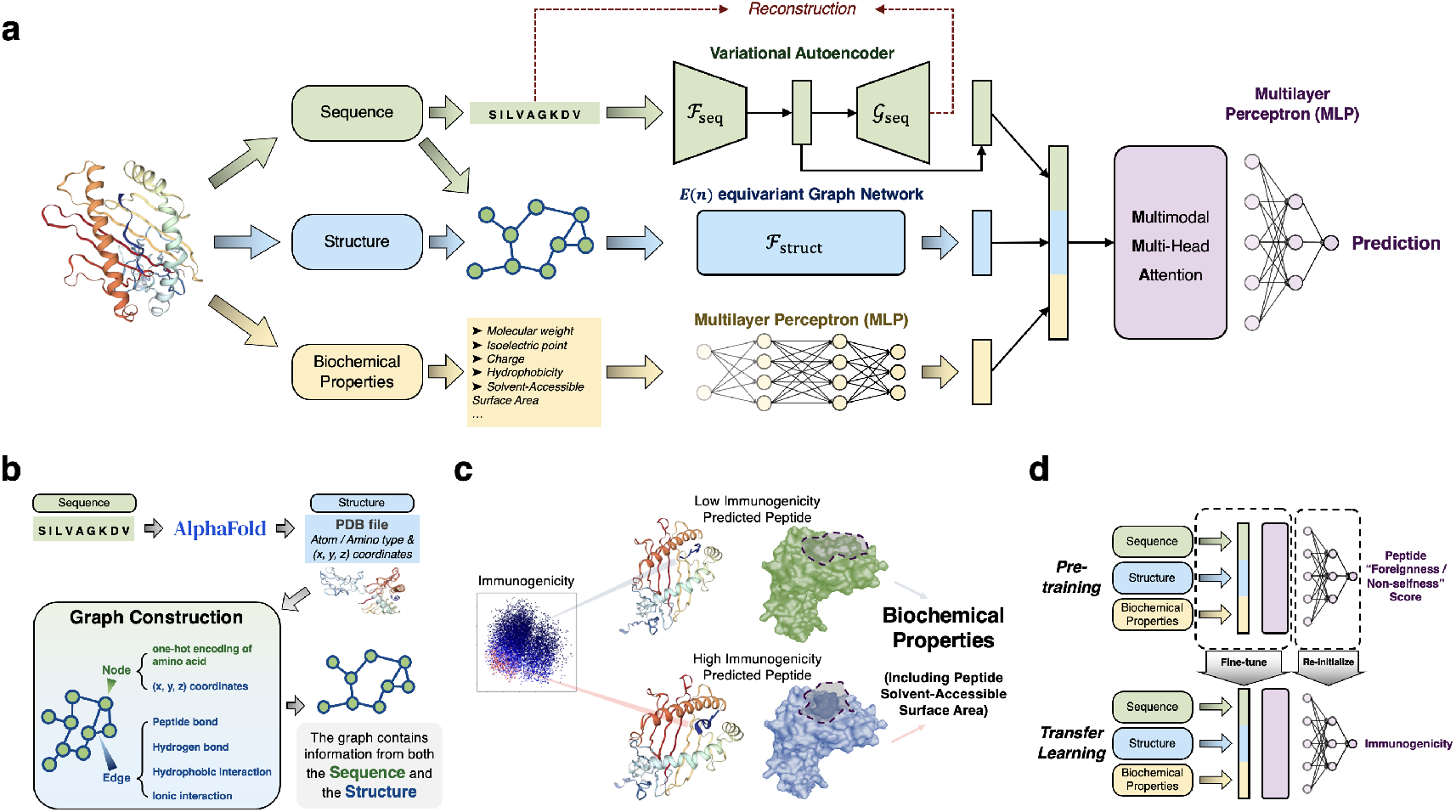
Overview of ImmunoStruct architecture and training. **(a)** ImmunoStruct separately encodes three modalities and uses the fused multimodal representations for downstream prediction. **(b)** Peptide-MHC graphs are constructed from AlphaFold2 output. Graph edges encode rich biochemical bond information within the peptide-MHC structure, while the nodes encode the 3D coordinates and the amino acid sequence. **(c)** Biochemical properties are computed from peptide-MHC structure and peptide sequence. **(d)** During transfer learning, the model is pre-trained to predict peptide foreignness and further tuned to predict immunogenicity.

MLP to produce a single scalar value for the prediction of immunogenicity. Technical details on data curation and processing are further described in Methods.

### Structure-sequence joint modeling

Peptide-MHC structures are first predicted from peptide and MHC sequences using a high-throughput AlphaFold2 pipeline where we jointly fold the peptide into the MHC-protein, thereby forming the peptide-MHC complex for each pair. These peptide-MHC structures contain both structural and sequence amino acid information, which are then converted into graphs that contain biochemical bonds of the peptide-MHC as the edges, as well as amino acid sequence and 3D positional information as the node features (Figure 1b). These graphs are then processed using a graph transformer module that also leverages *E*(*n*)-equivariant graph neural network (EGNN) [23] layers that are equivariant with respect to 3D rotation, translation, reflection, and permutation [24]. A multi-head self-attention mechanism enables it to capture spatial relationships and attend to learned pairwise interactions at all amino acid positions of the peptide-MHC [25]. This results in a 64-dimensional structural embedding, which are then integrated with the information processed from the other two modalities (Figure 1a). Since data are represented and processed as graphs, for simplicity, we will refer to this joint modeling as “Structure” in ablation studies.

### Biostructural interpretation module

To directly visualize the learned structural attention patterns for the immunogenicity of peptide-MHC, we implemented a structural attention visualization module. This enables interpretable overlays for individual amino acid residues, node-to-node amino acid interactions, and global attention-guided biostructural interpretations of the peptide-MHC surface (Supplementary Figure S1a-b). The module allows for fast extraction and processing of learned pairwise sequence-structure attention weights across all nodes of the peptide-MHC and can be easily integrated with the raw protein structure file of 3D atomic coordinates generated by AlphaFold2.

### Sequence modeling

We also explicitly encode peptide-MHC sequences by a variational autoen-coder (VAE) framework which is further processed by a multi-head self-attention layer [26]. The full sequences are obtained by concatenating the peptide, which is derived and processed from the full protein, and then concatenated to the end of the MHC sequence. We speculated that directly encoding the full peptide-MHC sequence information into a more compact and regularized low-dimensional space would enable learning additional sequence-based patterns and relationships across similar sequences and motifs. By encoding the full peptide-MHC sequences in this compact representation, we obtain a 32-dimensional latent space for each sample (Figure 1a) and reverse-map this to the input dimension to perform sequence reconstruction. This architecture choice is helpful for learning a compact and meaningful latent space that is semantically continuous across samples [26–28].

### Biochemical property modeling

Given the known biochemical involvement of interactions within the peptide-MHC-TCR synapse, and the importance of surface-area accessibility of the loaded peptide for immunogenicity [12, 17], we used the structural data for each peptide-MHC sample to calculate the solvent-accessible surface area (SASA) of the peptide-MHC to include as a component of the biochemical features (Figure 1c). The biochemical properties also included properties of each peptide, such as molecular weight, isoelectric point, charge, and hyrophobicity [7, 29]. The properties are then fed into the network as additional features (Figure 1a).

### Feature combination via multimodal multi-head attention

The 32-dimensional sequence embedding and the 64-dimensional structural embedding, along with the biochemical property embedding, are concatenated to form a 104-dimensional representation vector (Figure 1a). This multimodal representation undergoes further processing with a multi-head attention layer, which consists of eight self-attention heads applied directly to the multimodal representation vector. Attention visualization showed learned multimodal attention weights across the concatenated embedding, indicating information was attended to across each input modality (Supplementary Figure S1c).

### ImmunoStruct Training

#### Transfer learning

ImmunoStruct is trained using a transfer learning paradigm [30] (Figure 1d). After weight initialization, it is first pre-trained to predict *foreignness*, a measure that quantifies the similarity between a peptide and sequences of known pathogens – *foreign* to humans – which can be associated with immunogenicity [7, 31]. The score is computed using a multistate thermodynamic model as previously described [32]. After that, it is fine-tuned to predict immunogenicity, which is a binary experimental readout obtained from a T-cell recognition/activation assay that specifically detects recognition/activation of CD8+ T-cells to a given peptide-MHC. All modules are warm-started from the pre-trained weights, with the exception of the final MLP which is re-initialized to accommodate the change in task (Figure 1d). The transfer learning process leverages patterns between foreignness and epitope recognition [31, 33] to improve the performance on the main immunogenicity task that is different but related to the pre-training task. For both infectious disease and cancer models, graph data were generated from predicted AlphaFold2 structures augmented by sequences as the node features (Figure 1b), while biochemical property data were derived from sequence and structural surface features (Figure 1c). Details are further described in Methods.

#### Overview and Cancer mutant vs wildtype peptide-MHC contrastive learning

Given that the relative measured immune responses of infectious disease epitopes differs from that of cancer neoepitopes, we implemented a biologically motivated variation to our training strategy for application to cancer neoeptiope prediction. We propose a novel contrastive learning framework that organizes the latent space of ImmunoStruct such that cancer mutant peptides are embedded far from their wildtype counterparts if and only if the peptide-MHC is immunogenic (Figure 2a). This framework is based on the fundamental immunological concept of central and peripheral tolerance, whereby T cells that recognize self-derived peptide-MHCs are deleted or anergized (incapacitated) during thymic development [12, 34]. As a result, wildtype peptide-MHCs are typically immunologically silent, while mutated peptides from mutated cancer proteins can escape tolerance by altering TCR-facing residues and can elicit T cell responses if sufficiently distinct from their wildtype forms. In line with this, as well as previous work highlighting the differences between a wildtype and mutant peptide within the MHC binding groove [12], we reasoned that helping the model learn patterns related to these different multimodal representations may be more in line with the underlying immunology that potentiate the immune response. With respect to these representations, the main idea is that a cancer peptide-MHC should be embedded far away from its corresponding wildtype version if it is immunogenic, and close to it otherwise (Figure 2b). Details are further described in Methods.

**Figure 2:**
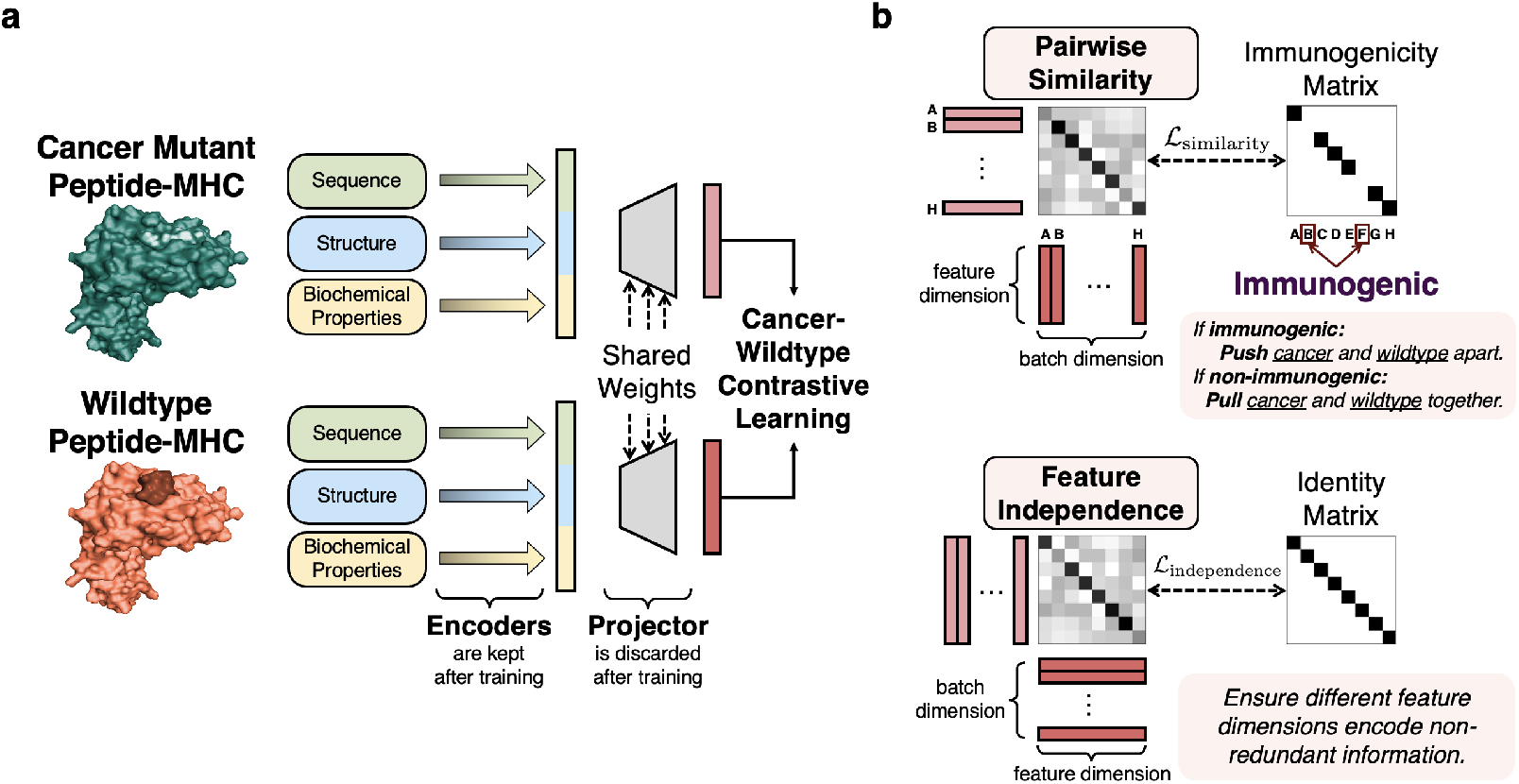
Cancer-wildtype contrastive learning leverages the wildtype counterpart to better organize the latent space. **(a)** ImmunoStruct extracts multimodal latent representations by integrating sequence, structure, and biochemical features. Cancer-wildtype contrastive learning optimizes these representations by contrasting cancer mutants with their wildtype counterparts. **(b)** Cancer-wildtype contrastive learning consists of two complementary components: the pairwise similarity component pulls or pushes latent representations depending on the immunogenicity of the peptide-MHC, and the feature independence component encourages different dimensions of the representations to encode diverse and distinct information.

### ImmunoStruct improves immunogenicity prediction on infectious disease epitopes and highlights positions important for prediction

We first evaluated ImmunoStruct for its ability to predict peptide-MHC immunogenicity on the IEDB dataset of infectious disease epitopes, and compared it with existing methods including Prime-2.0 [21], NetMHCpan [13], MHCnuggets [14], MHCflurry [15], BigMHC [16], and DeepNeo [22]. Performances are reported on AUROC, AUPRC and mean PPVn. AUROC and AUPRC are composite metrics that reflect the threshold-agnostic aggregated classification performance on {sensitivity, specificity} and {precision, recall, respectively}. Mean PPVn is a widely used and clinically important metric [13–16] that measures the precision of several top-ranked peptide-MHC candidates predicted in a setting similar to epitope ranking for clinical trial implementation.

For all metrics, ImmunoStruct established a new state of the art, with an AUROC of 0.882 *±* 0.005, an AUPRC of 0.696 *±* 0.020, and a mean PPVn of 0.514 *±* 0.020, as shown in Figure 3a. Notably, on the key measure of AUPRC, ImmunoStruct (0.696) largely outperformed PRIME2.1 (0.212), NetMHCpan (0.214), MHCnuggets (0.246), MHCflurry (0.260), DeepNeo (269), BigMHC-EL (0.319), and BigMHC-IM (0.462). The results can be found in Figure 3a and Supplementary Table S2.

**Figure 3:**
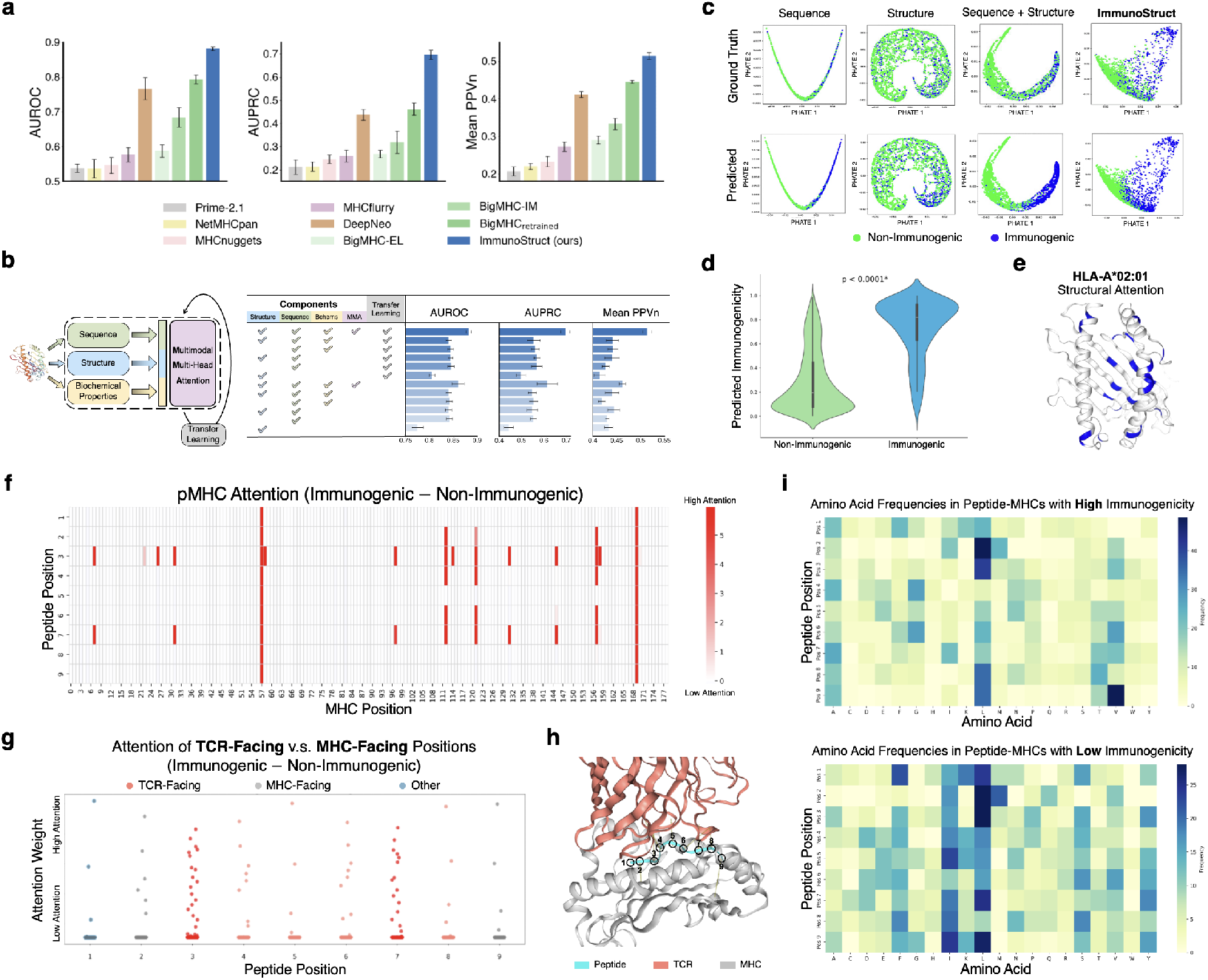
ImmunoStruct improves immunogenicity prediction on the **IEDB dataset** of infectious disease. **(a)** The performance of ImmunoStruct compared against existing methods. Mean value and standard deviation (AUROC, AUPRC) or standard error of the mean (Mean PPVn) from 5 independent experiments are displayed. For trainable models, results are reported on the test set after the models are trained on the non-overlapping training data. For pre-trained models with public API but without open-source code, evaluations are performed on the 5 test sets. **(b)** Ablation studies suggest that transfer learning is helpful and that combining all components with multimodal multi-head attention leads to the best performance. **(c)** Embedding visualizations between ground truth (top) and predicted (bottom) immunogenicity scores suggests that the ImmunoStruct model learns a structured latent space that can well distinguish immunogenic and non-immunogenic samples by fusing multimodal information (sequence, structure, and biochemical properties). **(d)** Immunogenicity scores predicted by ImmunoStruct over all samples in the test set show a clear separation between immunogenic and non-immmunogenic peptide-MHCs. **(e)** The structural attention visualizations shows the high-attention locations of the MHC protein structure. **(f)** Learned immunogenicity-enhanced attention between peptide and MHC positions. **(g)** Averaged attention across peptide positions. The TCR-facing positions 3 and 7 show high attention. **(h)** The structural visualization shows that peptide positions 2 and 9 are MHC-facing and peptide positions 3 through 8 are TCR-facing. More views available in Supplementary Figure S2. **(i)** Heatmap of amino acid frequencies in high immunogenicity (top) versus low immunogenicity (bottom) peptides. **(e)***−***(i)** are generated on allele HLA*A:0201.

To verify that all input modalities are beneficial and synergistically improve model performance, we conducted an ablation experiment (Figure 3b and Supplementary Table S1). The ImmunoStruct model that takes in all input modalities outperformed all of its ablated counterparts, and showed a synergistic pattern upon including all components and modalities. This study also demonstrated that transfer learning is generally beneficial compared to training from scratch.

The same phenomenon can be observed qualitatively as well. By visualizing the latent space embeddings using PHATE [35], we observed that the latent space of ImmunoStruct with all three modalities exhibited the clearest organization when comparing the true vs predicted labels (Figure 3c). These visualizations indeed show an increasingly organized embedding space of peptide-MHC immunogenicity, colored by true immunogenicity (top row) vs bottom row (predicted). This embedding organization becomes most pronounced for the ImmunoStruct model.

We next leveraged the ImmunnoStruct EGNN attention layer to assess which positions on both the peptide and MHC were important for prediction. For easy comparison of protein index positions, we performed this analysis on a single allele (HLA*A:0201). We first show strong separation in ImmunoStruct prediction scores between non-immunogenic and immunogenic samples (Figure 3d). Next, we extracted the learned structural attention weights and overlayed them onto the MHC structure, showing the high-attention

MHC enrichment throughout the *α*1 and *α*2 peptide-binding cleft domains, and the N-terminal extracellular domain (Figure 3e).

Interestingly, when assessing peptide level structural attention weights (Figure 3f-g), we observed enrichment in learned attention values at TCR-facing positions – particularly positions 3, 4, 6 and 7 of the peptide – as opposed to MHC-facing positions (e.g., 2 and 9) which would be more related to peptide-MHC binding rather than immunogenicity [36] (Figure 3h). Upon assessing these positions more closely, we found that immunogenic samples display less diversity at these TCR-facing positions (Figure 3i). Visualization of these positions on a known example peptide-MHC-TCR structure also show protrusion of these peptide positions towards the TCR, which aligns with the critical role of TCR interactions in the immunogenic response and may suggest increased sequence-structural focus towards these positions in immunogenic samples (Figure 3h). We also observed differences in amino acid frequencies between immunogenic and non-immunogenic peptides (Figure 3i), as well as a higher enrichment of alanine at position 7 and preference of leucine at position 3. The TCR-facing and MHC-facing positions are illustrated in Figure 3h, with more views available in Supplementary Figure S2. Together, these results suggest ImmunoStruct could be leveraged to pinpoint certain peptide-MHC positions that are important for immunogenicity prediction.

### ImmunoStruct improves immunogenicity prediction on human cancer neoepitopes

The clinical potential of therapeutic cancer neoepitiope-based vaccines has been increasing in recent years [1– 3], so we next sought to explore the prediction performance of ImmunoStruct in the setting of cancer neoepitope immunogenicity. This task may improve prioritization of the most immunogenic peptides for epitope-based vaccine design, which is restricted to a limited number of peptide candidates. Missense mutations to a protein can lead to presentation of a peptide that only differs by one amino acid from its wildtype form, which may make distinguishing immunogenic neoepitopes challenging [12, 37].

Upon extending ImmunoStruct to this data setting, we observed superior performance compared to existing methods (Figure 4) despite a relatively smaller training set than some other competitors. ImmunoStruct was able to outperform existing methods on AUROC, AUPRC and mean PPVn (see Figure 4a and Supplementary Table S3). We also showed through an ablation study that the cancer-wildtype contrastive learning framework and the transfer learning paradigm both additively improved ImmunoStruct performance (Figure 4b).

**Figure 4:**
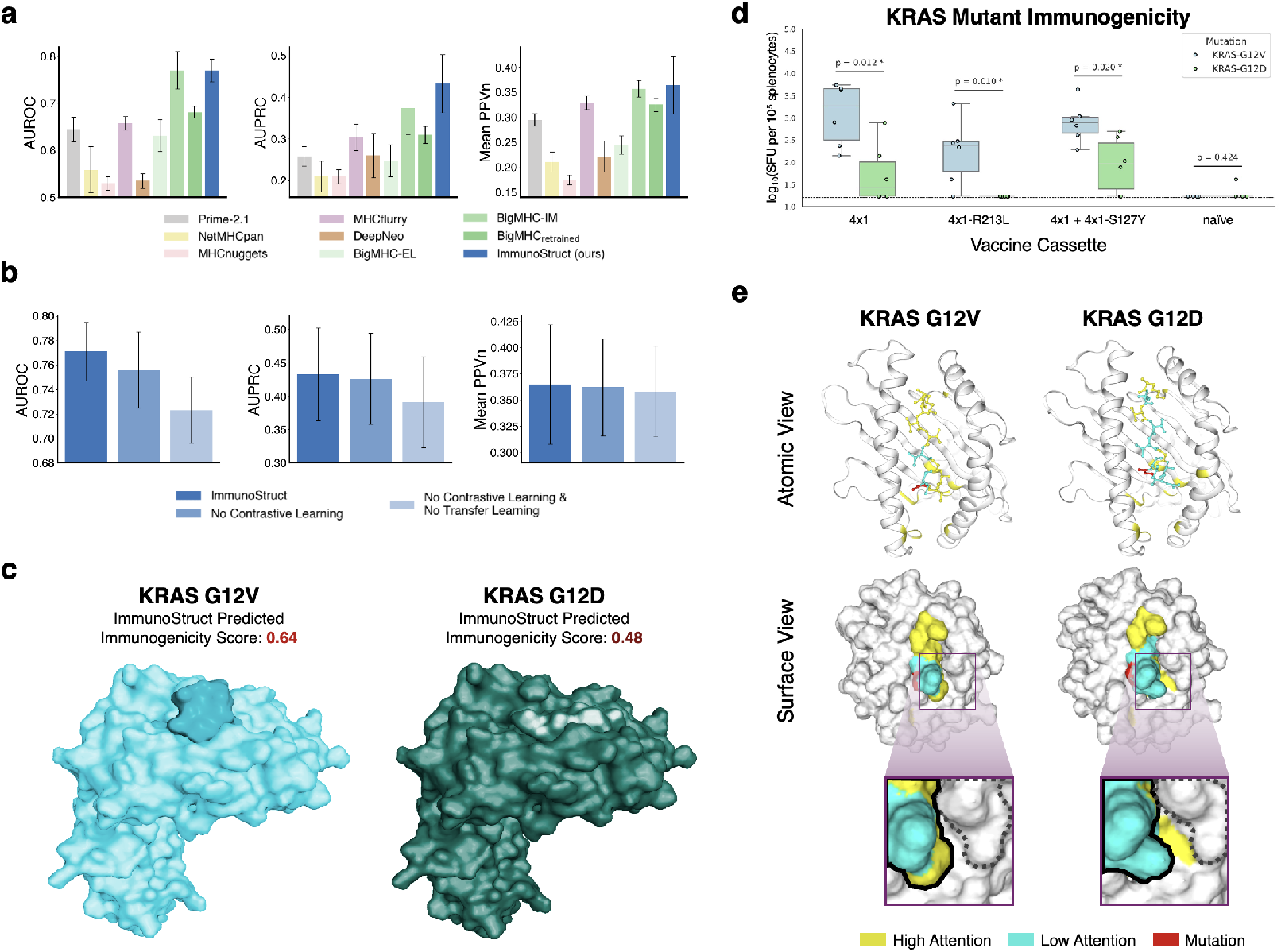
ImmunoStruct improves immunogenicity prediction on **cancer neoepitopes. (a)** The performance of ImmunoStruct compared against existing methods. Mean value and standard deviation (AUROC, AUPRC) or standard error of the mean (Mean PPVn) from 5 independent experiments are displayed. For trainable models, results are reported on the test set after the models are trained on the non-overlapping training data. For pre-trained models with public API but without open-source code, evaluations are performed on the 5 test sets. **(b)** Ablation studies suggest that transfer learning and cancer-wildtype contrastive learning are both helpful. **(c)** Protein molecular surface visualization of KRAS mutants with ImmunosStruct Immunogenicity Scores. **(d)** Experimental KRAS Mutant Isoform Immunogenicity for three distinct vaccine constructs containing either the G12V or G12D isoform or a naive negative control **(e)** KRAS mutant peptide-MHC visualization with ImmunoStruct attention overlay. Solid lines outline the peptide; dotted lines outline nearby MHC protein residues.

Given the structural subtleties in cancer neoantigen peptide-MHC immunogenicity particularly for oncogene mutant isoforms, we next sought to identify and explore the ability of ImmunoStruct to properly capture these differences and result in appropriate predictions. To that end, we first ran ImmunoStruct on the KRAS-G12V and KRAS-G12D peptide-MHC (HLA-A*11:01) mutants as a test case analysis, resulting in increased immunogenicity predictions for the G12V mutant compared to the G12D mutant (0.64 vs 0.48), as visualized in Figure 4c. Indeed, when experimentally measured within the clinical neoantigen vaccine context (NCT03953235 [38]), the KRAS-G12V mutant displayed increased immmunogenicty compared to the KRAS-G12D isoform across three separate vaccine cassettes (p *<* 0.05, Figure 4d). These results suggest the abilitiy of ImmunoStruct to capture subtle peptide-MHC multimodal differences resulting from a single isoform amino acid change from the same protein.

More importantly, we want to explore which parts of the peptide-MHC structure contribute the most to the prediction of immunogenicity and visually compare the differences between the two isoforms. We extracted the learned ImmunoStruct attention weights from the structural multi-head attention layers. To date and to our knowledge, there are no structural graph attention-based deep learning methods from wide-scale peptide-MHC structure data for immunogenic biostructural interpretation. We therefore built a attention visualization module that helps highlight critical regions of the peptide-MHC that influence the predicted immunogenicity, providing insights into structural determinants of epitope recognition. Interestingly, when extracting the learned attention weights, we observed a region of the peptide-MHC groove surface that showed differential sterics between the two isoforms which aligned with the learned attention weight at those positions (Figure 4e). This suggested that towards the terminal end of the KRAS G12V peptide presented, a differential peptide-MHC interaction may contribute in part to a higher immunogenic response, as has been observed in the context of other peptide-MHCs [39]. Additional example cases of structural attention from the visualization module are shown in Supplementary Figure S1.

### Experimental validation on SARS-CoV-2 epitopes and clinical cancer survival prediction

Given the importance of further validation for an immunogenicity prediction model, we sought to validate our predictions with ELISpot IFNg measurements that we generated *in vitro* for a set of peptides in combination with one of five HLA alleles. ImmunoStruct successfully predicted 15 of 19 immunogenicities (AUROC = 0.780, AUPRC = 0.941, and accuracy = 79.0%) (Figure 5a). For each allele, one plate is shown to visualize IFNg production as an example plate for a given peptide-MHC (Figure 5b), further described in Methods.

**Figure 5:**
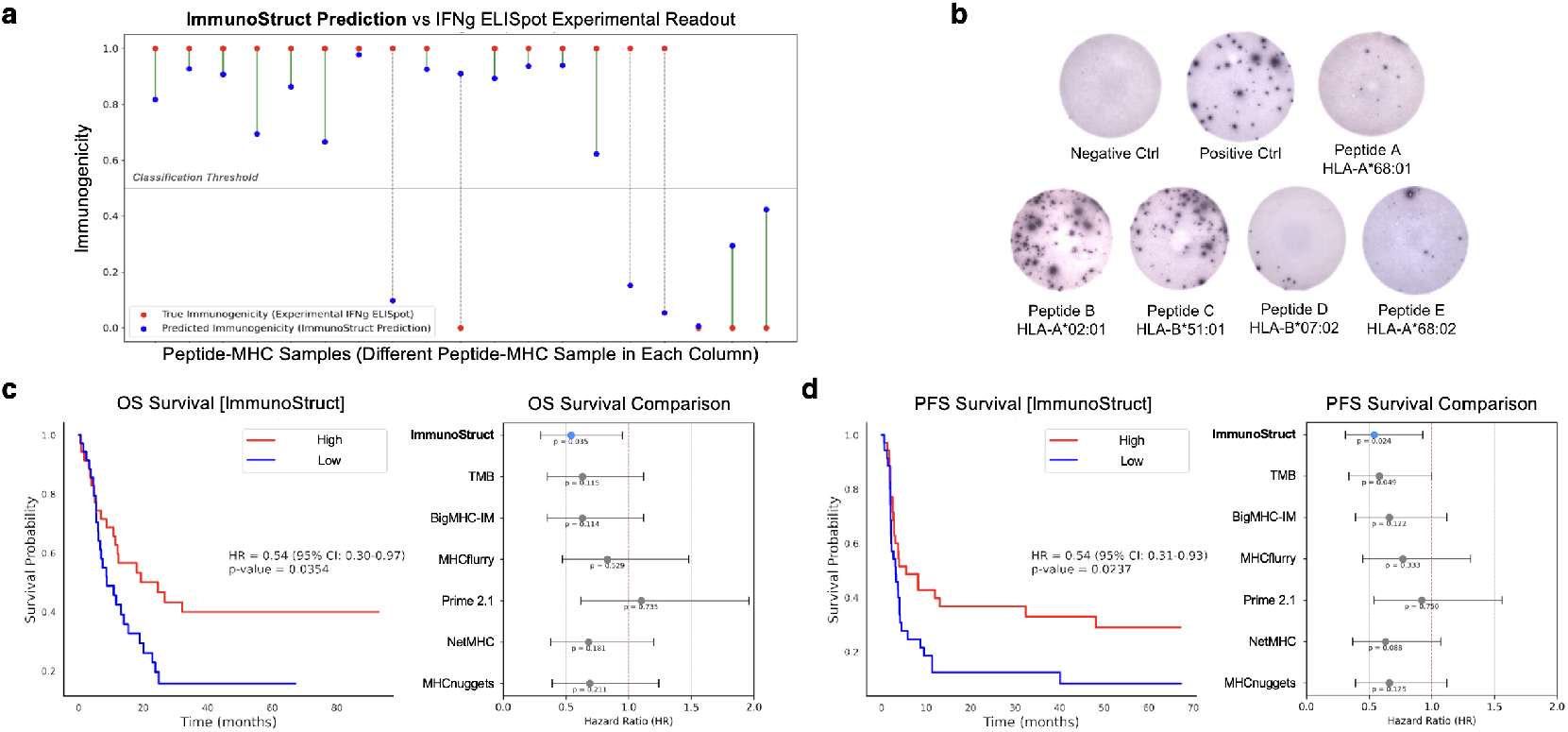
Experimental validation of ImmunoStruct predictions. **(a)** Lollipop plot where each column represents a unique epitope-MHC pair. Red dots indicate the experimentally measured immunogenicity with IFNg ELISpot and blue dots indicate the predicted immunogenicity from ImmunoStruct. Vertical lines connect paired values for each sample, with green lines for correct predictions and dashed gray lines for incorrect predictions. **(b)** ELISpot plate images, with each plate corresponding to a validated allele. **(c)**–**(d)** Kaplan–Meier survival analyses stratified by the median count of immunogenic peptide–MHC. ImmunoStruct-predicted immunogenicity demonstrates strong association with survival (left panel), outperforming existing methods (right panel). Overall survival (OS) is shown in **(c)**, and progression-free survival (PFS) is shown in **(d)**. Horizontal error bars indicate 95% confidence intervals. Survival curves for all methods compared can be found in Supplementary Figure S3.

To extend the utility of ImmunoStruct to the clinical cancer setting, we validated ImmunoStruct in a cohort of cancer patients with available peptide-MHC data, comprising a variety of different tumor types and immunotherapeutic treatments [37]. In comparison to tumor-mutational burden, we found ImmunoStruct predictions were able to stratify patient survival more effectively for both overall survival and progression-free survival without any re-training (HR = 0.54 [0.30-0.97] and HR = 0.54 [0.31-0.93], respectively, Figure 5c-d). We also show comparisons with existing methods, indicating improved performance on peptide-MHC-based prediction of cancer patient overall survival and progression free survival (Figure 5c-d, and Supplementary Figure S3). Together, these results suggest the ability of Immunostruct to capture clinically relevant immunogenic peptide-MHCs that associate with survival in the context of cancer immunotherapy.

## Discussion

The ImmunoStruct model is the first jointly trained structure-sequence model to use large-scale multiallelic structural and biochemical peptide-MHC data for an application to the immunogenicity prediction task, across both infectious disease and cancer neoepitopes. Immunostruct makes novel use of graph transformer, MLP and VAE neural networks for different modalities, culminating in a multimodal architecture capable of synergistically integrating three distinct data modalities for this complex task. Second, within the graph transformer, the multi-head self-attention mechanism provides biostructural insights for immunogenic peptide-MHC features and the ability to visualize regions of interest in a peptide-MHC complex. Third, the new contrastive loss approach for cancer neoepitopes improves prediction performance by embedding wildtype and mutant peptide sequences for immunogenic peptide-MHCs far in spite of only differing by a single amino acid. Lastly, we also provide a curated multimodal dataset containing structural, sequence, and biochemical property information for each labeled peptide-MHC sample.

When assessing the design steps and modalities of ImmunoStruct, each component was biologically inspired, aiming to extract unique information from each peptide-MHC data type. For example, the structure embedding encoded from the graph transformer module allowed for precise encoding of spatial relationships that are relevant for TCR binding to the peptide-MHC surface [12]. The peptide-MHC surface characteristics where further accounted for by the solvent-accessible surface area feature included in the model. Such information may otherwise be overlooked by sequence-only models. The pre-training paradigm on peptide foreignness was also inspired by the concept of T-cell self-tolerance mechanisms and higher observed immunogenicity to foreign peptides [7, 31, 33], showing an added increase in performance. For structures we generated from Alphafold, our goal was to determine whether there would be a net change in immunogenicity prediction performance with the added structural information. Interestingly, we learned through our ablation studies that inclusions of all components together with our transfer learning paradigm, synergistically, rather than additively, improved performance. This may suggest that there exist non-linear combinations of learned downstream feature vectors that contain meaningful information for future exploration of immunogenicity prediction.

When specifically exploring cancer neoepitopes, we added a conditional contrastive loss to ensure that the mutant peptide will be embedded far from its wildtype counterpart only if the mutation results in immunogenicity. This helped force the model to learn how to differentiate each pair and capture subtle differences in peptide-MHC variants. We applied this to a case study for the known and highly prevalent cancer oncogene KRAS, which is a clinical target across several solid tumor types [38]. Specifically, we saw that when leveraging experimentally validated KRAS-G12V vs KRAS-G12D with in-vivo samples, we were able to leverage our attention-based visualization module to highlight subtle peptide-MHC spatial proximities that differed within the peptide-binding groove between each complex. Interestingly, the attention difference captured was at the C-terminal end of the peptide rather than at the mutation itself, suggesting a conformational peptide-MHC effect rather than the mutational effect at the sequence level itself. Interactions such as these observed from ImmunoStruct may encourage interesting avenues of machine-guided future exploration in the domain of cancer neoantigen vaccine design as well as TCR design. Additionally, given these subtle conformational differences may exist across different samples and MHC alleles, a wide-scale generation and integration of molecular dynamics peptide-MHC data may also provide an interesting future direction of our work.

Despite these contributions and interesting avenues of future exploration, the study has limitations. First, as a general limitation in the field of immunogenicity prediction, the peptide-MHC data used for training relies on experimental labels, which are inherently variable to some degree due to differences in assay protocols, cell lines, and experimental designs. Second, while TCR information would allow more information for prediction, the astronomical diversity of the human TCR repertoire across peptide-MHCs limits training data availability. Third, ImmunoStruct was trained on only the 27 most common HLA allele protein sequences [40], and therefore further analysis would be required to explore its application to rare alleles with low training data. Finally, the application to clinical neoeptiope survival consisted of a small sample size and was not homogenized with regard to a single immunotherapy type. However, this also highlights the need for additional publicly available clinical-genomic immunotherapy datasets for validation of tools within this field.

Together, ImmunoStruct provides a multi-modal structural, sequential, and biochemical approach that is meaningfully able to jointly model protein structure and sequence for the challenging task of immunogenicity prediction. With the advent of mRNA cancer neoantigen vaccines, as well as immunogenicity bio-therapeutic toxicity challenges, this work has applications to not only epitope-based vaccines, but also protein-based therapeutic discovery. More generally, it may also provide a framework for protein-complex property prediction and protein design goals that may benefit from integrating biochemical and predicted structural data.

## Methods

### ImmunoStruct

#### Structure-sequence modeling with multimodal graph transformer

To create the peptide-MHC structure dataset which also contain sequence features, we first use AlphaFold2 [20] to generate PDB files containing the folded protein structures from the peptide-MHC sequences. For each peptide-MHC sequence, the protein structures are represented as the amino acid residue sequence information and their respective (*x, y, z*) coordinates in the 3D space. First, we trim the structure to take in consideration only the extracellular domain of the peptide-MHC. We then construct a graph 𝒢= (𝒱, ℰ) where the nodes 𝒱= *{v*_1_, *v*_2_, …, *v*_*n*_*}* are the one-hot encoding of the amino acid residues concatenated to their (*x, y, z*) coordinates, whereas the edges ℰ= *{e*_1_, *e*_2_, …, *e*_*m*_*}* are binary existence indicators of peptide bonds, hydrogen bonds, hydrophobic interactions or ionic interactions between two vertices. Specifically, a node *v*_*i*_ is represented as a 1 *×* 23 vector [AA_1_, …, AA_20_, *x, y, z*]^*⊤*^ where AA_[0:20]_ one-hot encodes the amino acid residue. An edge *e*_*ij*_ is defined as 1 if there exists any aforementioned bond or interaction between *v*_*i*_ and *v*_*j*_, and as 0 otherwise.

When modeling the peptide-MHC structures, ImmunoStruct leverages a graph transformer module with *E*(*n*)-equivariant layers (EGNN) [23] and multi-head self-attention (later described in section “Multimodal data integration strategy”) across all node-sequence and spatial information of the peptide-MHC structure. The EGNN layers enable powerful processing of data represented as graphs, where nodes correspond to amino acid residues and edges represent the interactions among residues. EGNN layers further amend the graph neural network by ensuring equivariance to 3D coordinates. For this task these equivariant layers are suitable because they are equivariant to rotations, translations, reflections and permutations so the arbitrary positions and orientations from the AlphaFold 3D coordinates would otherwise be properly handled.

The propagation rules from layer *l* to *l* + 1 is described in Eqn (1). *h* is the node embedding, *x* is the coordinate embedding, *a* is the edge attributes, and the *ϕ*’s are learnable operations. Following the official implementation [23], *C* is set to be 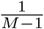, where *M* is 3 for the 3D coordinates.

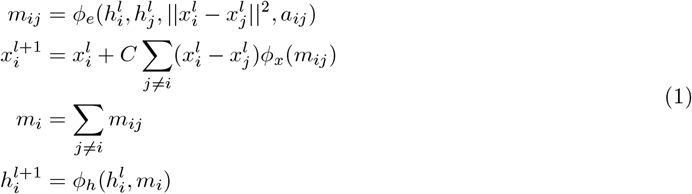

In our case, 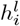 is the embedding of residue *v*_*i*_ at layer 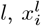 are the coordinate embeddings from spatial coordinates (*x, y, z*), and *a*_*ij*_ are the edge attributes as previously described.

Importantly, the combination of multi-head self-attention and graph processing layers ultimately enable the model to capture the spatial and pairwise relationships across the peptide-MHC, which are cooperatively recognized by a complementary TCR structure within the immunogenic synapse.

#### Sequence modeling with a variational autoencoder

The peptide-MHC sequences are encoded via a variational autoencoder (VAE) [26]. The VAE learns a continuous and structured latent space by introducing a probabilistic framework. It uses an encoder *g*_*ϕ*_ and a decoder *f*_*θ*_ to encode the input *x* to a latent vector *z* and reconstruct it back to 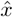 which approximates the input. In a standard autoencoder, the process is described in Eqn (2) and the model is optimized by minimizing the L2-distance between the real and reconstructed input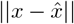.

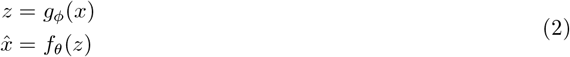

VAE introduce a probabilistic framework by separately modeling the mean and variance of the data, as described in Eqn (3), where *⊙* stands for element-wise product. Besides minimizing the L2-distance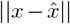, it also minimizes the KL divergence between the latent embedding *z* and a standard normal distribution 𝒩 (0, *I*). This allows the VAE to learn a semantically continuous latent space, since latent embedding *z* are reconstructed to approximate the same input despite small perturbations *σ ⊙ ϵ*. For more details on the optimization, please refer to section “ImmunoStruct optimization objective”.

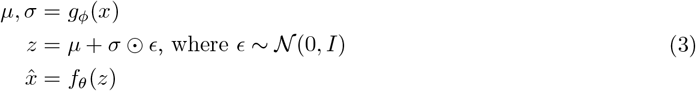

In our implementation, the VAE module projects the higher-dimensional sequence data into a lower 32-dimensional latent space. Because of the semantic continuity property, similar peptide-MHC sequences are embedded closely. This is beneficial to the immunogenicity prediction task as sequence-based motifs are known to associate with immunogenicity. Lastly, the semantically continuous latent space is easily amendable to downstream data integration with other data modalities, namely structure and biochemical properties.

#### Biochemical property modeling with a multi-layer perceptron

A multilayer perceptron (MLP) that consists of two linear layers with ReLU activation and a dropout layer is used to extract an embedding vector from the raw biochemical property values. This light-weight module allows the learning of higher-order non-linear combinations of these properties that serve as helpful features that are biochemically derived for the downstream task.

#### Transfer Learning over peptide *foreignness*

Before training the model to perform the binary classification task of immunogenicity, our model is first trained to predict *foreignness scores* as defined by [7, 31, 33]. The foreignness can be seen as a measure of how similar a sequence is to sequences of known pathogens and was shown to be related to immunogenicity.

As defined by [33, 41] foreignness (*R*) follows a thermodynamic model as:

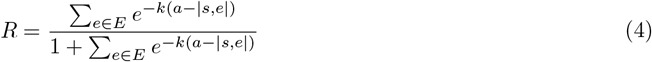

We used the same parameters as [41] Where *E* is the dataset of pathogen sequences, *k* = 4.87, *a* = 26 and |*s, e*| defines how similar in sequence *s* is from *e* using computed alignment scores.

This score enabled us to first train a regressor and then perform the final task. The ablation studies (Figure 3a) showed that this transfer learning method improved performance.

#### Multimodal data integration strategy

After encoding of each data-modality in latent space, with *h*_sequence_ ∈ ℝ^32^, *h*_structure_ ∈ ℝ^64^, *h*_bchems_ ∈ ℝ^8^, the modality-specific embeddings are concatenated into a single input vector *X* = *h*_sequence_ ∥*h*_structure_ ∥*h*_bchems_, *X* ∈ ℝ^104^. This concatenation aligns the features from heterogeneous sources into a single common embedding vector, ensuring that the downstream layers can jointly input the information from each modality. This vector is then processed using a multi-head self-attention mechanism [25] directly on the latent vector, which is a novel application to this domain.

The standard scaled-dot product attention is performed as follows. We first compute three vectors, namely query *Q*, key *K* and value *V* from the input vector *X* by matrix multiplications *Q* = *W*^*Q*^*X, K* = *W*^*K*^*X, V* = *W*^*V*^ *X*, where the *W* ‘s are separate learnable matrices. The attention is given by Eqn (5), where *d*_*k*_ is the dimension of the key vector.

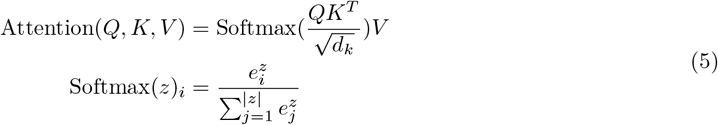

Multi-head attention [25] is an extension to the scaled-dot product attention. Instead of performing a single attention, it project the queries, keys and values *h* times with different, learned linear projections (Eqn 6). Multi-head attention captures diverse aspects of relationships within a sequence by allowing each “head” to focus on different parts of the data simultaneously.

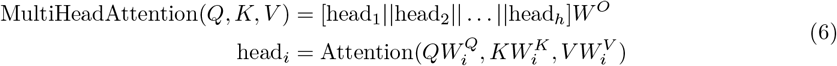

In our implementation, we used a multi-head attention mechanism that encompasses 8 attention heads to dynamically compute attention weights over the combined multimodal representation. This enables the model to selectively focus on the most informative features for a given sample’s prediction, rather than trivially treat the different modalities as if they were equivalent. For instance, in cases where structural nuances play a critical role such as when differentiating subtle peptide-MHC conformations which the peptide-MHC sequences alone would not inherently capture, higher attention weights can be allocated to the structural representations. Furthermore, each attention head can capture distinct cross-modal relationships. This mechanism also inherently provides a form of regularization by allowing the network to down-weight less informative or noisy features from any given modality. This is beneficial in contexts like ours, where peptide-MHC labels and data quality can vary depending on its source and experimental protocols.

#### Visualization of structural peptide-MHC attention

A single self-attention layer (as described in Eqn 5) was applied directly on the peptide-MHC graphs during training of the EGNN module to capture sequence-structural relationships directly from the peptide-MHC graph. After training, attention weights were extracted in the form of the attention matrix for each peptide–MHC (peptide-MHC) complex. We then isolated the submatrix corresponding to interactions between the peptide and MHC node features, which also contained the sequence information. As the raw attention values are typically very small, we applied a logit transformation across values to enhance subtle differences learned across the peptide. These were then averaged across samples and compared between immunogenic and non-immunogenic complexes across positions of the peptide.

#### ImmunoStruct optimization objective

During training, ImmunoStruct is simultaneously optimized by a prediction loss *ℒ*_pred_, a protein sequence reconstruction loss *ℒ*_recon_, and a normal distribution regularization on the latent space *ℒ*_normal_. The loss function is given by Eqn (7), where the loss components are weighted by their respective coefficients *λ*_pred_, *λ*_recon_, *λ*_normal_.

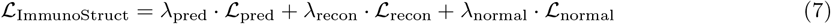

The prediction loss *ℒ*_pred_ guides the model to predict the target accurately, while the reconstruction loss *ℒ*_recon_ ensures that the learned representations carry sufficient information to faithfully reconstruct the input [27, 28, 42–44]. Additionally, the normal distribution regularization *ℒ*_normal_ encourages the latent space of the VAE to approximate a standard normal distribution with zero mean and unit variance [26].

The loss components are shown in Eqn (8), (9), (10) and (11). 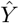 and *Y* are, respectively, the set of predicted and ground truth values to be predicted (foreignness or immunogenicity). 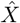 and *X* are, respectively, the set of predicted and ground truth protein sequences. 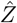 is the set of latent embedding vectors at the latent space of the VAE, while *Z* follows the normal distribution *𝒩* (0, **I**). *N* is the batch size and 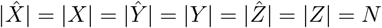.

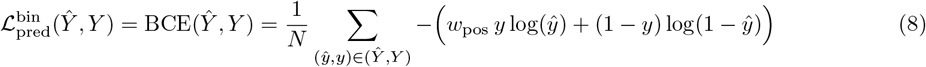

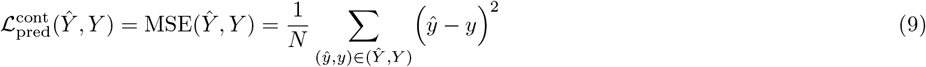

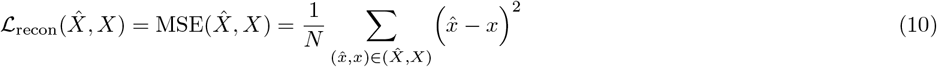

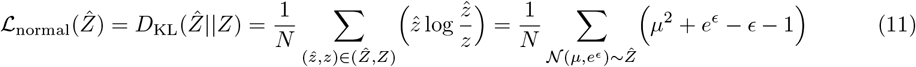

It should be noted that the prediction loss comes in two alternative forms: When the prediction target is binary (i.e., for immunogenicity), it uses the binary cross entropy loss (Eqn (8)); When the prediction target is continuous (i.e., for foreignness), it uses the mean squared error loss (Eqn (9)). The binary cross entropy loss is implemented using BCEWithLogitsLoss in PyTorch [45], where *w*_pos_ is the positive class weighting coefficient. Following the official recommendation from PyTorch, we set *w*_pos_ as the ratio of negative to positive sample counts.

For numerical stability, the normal distribution regularization is implemented using the reparameterization trick [26], where two separate linear layers in the neural network are respectively used to produce the mean *µ* and log variance *ϵ* of the latent representations. Then, the KL divergence can be computed using summations, subtractions, and exponents, without involving divisions (Eqn (11)).

#### Cancer-wildtype contrastive learning

When training the cancer Immunostruct model, multimodal inputs derived from both the mutant and wildtypes peptide-MHCs are leveraged. In practice, a light-weight projector network first projects the multimodal embeddings to a 128-dimensional latent space where cancer-wildtype contrastive learning is performed. The projector, known to be critical for contrastive learning [46, 47], is only used to indirectly influence the upstream multimodal embeddings and is discarded after training is complete.

As described in Eqn (12), The optimization objective is a cancer-wildtype contrastive loss *ℒ*_CW_ (Eqn (13)) scaled by coefficient *λ*_CW_ and added to the original loss function *ℒ*_ImmunoStruct_. The cancer-wildtype contrastive loss *ℒ*_CW_ consists of two main components, specifically an immunogenicity-aware pairwise similarity loss *ℒ*_similarity_ (Eqn (14)) and a feature independence loss *ℒ*_independence_ (Eqn (15)).

The immunogenicity-aware pairwise similarity loss *ℒ*_similarity_ organizes the latent embeddings across all cancer-wildtype pairs, pulling together non-immunogenic pairs and pushing apart immunogenic pairs. This is to enforce large embedding based distances for mutations that are experimentally observed to result in truly immunogenic neoantigens. The feature independence loss *ℒ*_independence_ encourages different feature dimensions to encode mutually non-redundant information and therefore mitigates dimensional collapse. The cancer embedding *Z*_C_ and the corresponding wildtype embedding *Z*_W_ share the dimension *N × D*, where *N* is the batch size and *D* is the feature dimensionality. The immunogenicity vector 𝟙^C^ is a binary vector of size *N*, where its element takes a value of 1 if the corresponding peptide-MHC is immunogenic and 0 otherwise.

While the feature independence loss is inspired by a prior work [47], the immunogenicity-aware pairwise similarity loss is carefully designed to take the advantage of the unique information from the cancer-wildtype pairs, and is our original contribution.

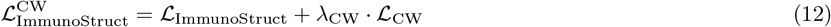

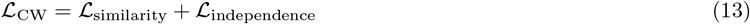

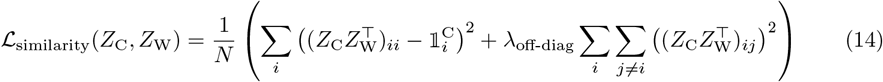

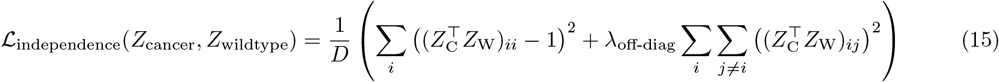

### Hyperparameters and training details

All experiments were performed on a 40-GB NVIDIA A100 GPU, and repeated under 5 random seeds. For all experiments, we used a batch size *N* of 128, a learning rate of 1 *×* 10^*−*3^ for pre-training and 1 *×* 10^*−*4^ for finetuning. The models were optimized with the AdamW [48] optimizer with a weight decay of 1 *×* 10^*−*6^. During pre-training, the learning rate was kept constant for 40 epochs. During finetuning, the learning rate was warmed up linearly for 10 epochs, followed by a cosine decay for 30 epochs [49]. For experiments following the transfer learning paradigm, both pre-training and finetuning are performed. For experiments marked as “no transfer learning”, the pre-training stage is skipped and the model was trained from scratch with finetuning.

The coefficients in the loss function (Eqn (7)) were empirically set to the following values: *λ*_pred_ = 2.0, *λ*_recon_ = 0.5 and *λ*_normal_ = 0.5. In cancer-wildtype contrastive learning (Eqn (12)), the coefficient was empirically set to *λ*_CW_ = 0.01.

### Multimodal Dataset for Immunogenicity Prediction

Due to the scarcity of publicly available peptide-MHC structural data, we implemented a high-throughput AlphaFold2 pipeline to generate the largest peptide-MHC structural dataset to our knowledge. This dataset ultimately comprised of approximately 27,000 entries with paired experimental immunogenicity labels.

#### Peptide-MHC Sequence Data

In order to curate a dataset that would provide sufficient epitope sequence training data across alleles, we downloaded data from the immune epitope database (IEDB) [18]. Peptide sequences were filtered to several criteria: (1) human sequences, (2) T-Cell experimental assay data (ELISpot, Tetramer staining), and MHC class-I restriction. To avoid training across rare human alleles with small and often only negative sample sizes, we curated a dataset for the top 27 most common alleles in the human population (*∼* 97% human population coverage) as previously described [40, 50]. The cancer peptide-MHC sequence data were obtained as previously described and filtered to the same allele set [18, 37, 40, 50]. In total, this provided a dataset of 27,000 peptide-MHC pairs with known positive or negative experimental epitope labels.

#### Peptide-MHC AlphaFold2 Structure Data

The AlphaFold2 pipeline was implemented as proposed previously [20]. To run our high-throughout dataset of peptide-MHC sequence combinations, we made use of ColabFold [51], a wrapper on top of AlphaFold2 that enables highly customizable and easily useable protein folding operations. With ColabFold, we were able to generate a protein folding from peptide-MHC pairs with only a few lines of code and a shared set of configurations. This enabled us to distribute our protein generation across multiple hosts, drastically reducing the amount of time taken from several weeks to a couple of days. After the proteins were generated, we applied post processing to the structures to convert them to a deep learning model readable graph format containing hydrogen-bond interactions, ionic interactions, and hydrophobic interactions, as well as amino acid node features and their 3D positional information.

#### Biochemical Properties

Reasoning the potential benefit of including biochemical properties as additional features in our prediction task, we integrated biochemical properties into the ImmunoStruct prediction. The peptides package [29] was used to calculate all possible 84 biochemical properties for each peptide. The top 10 biochemical properties with the highest standard deviation were selected in addition to the peptide-MHC solvent-accessible surface area score (SASA), computed as previously described [52]. These biochemical/biostructural properties were then grouped into two meta-property features (Bchem1 and Bchem2) to be used as additional features during training.

#### Peptide-MHC *in vitro* Immunogenicity Validation

IFN-gamma-based enzyme-linked immunospot (ELISpot) assays were used to validate a set of SARS-CoV-2 epitopes. ELISpot assays provide a standardized measurement of T cell responses [53, 54]. Previously frozen human peripheral blood mononuclear cells (PBMCs) were thawed rapidly to 37^*°*^C and incubated overnight (Millipore, Massachusetts, U.S.A.). PBMCs were plated at 400,000 cells/well and co-incubated with 10 *µ*g/mL of peptide for 48 hours at 37^*°*^C. ELISpot plates were analyzed using CTL Immunospot. Positive controls were PBMCs plated at 200,000 cells/well and co-incubated with Spike Glycoprotien peptide megapool at 1 *µ*g/mL (JPT Peptide Technologies, Berlin, Germany). Negative controls were unstimulated PBMCs plated at 200,000 cells/well. Positive T cell responses were defined by greater or equal to the mean of negative controls + 3. Three additional true negative peptide-MHC samples validated through similar methods were identified through literature review to improve data balance [55].

## Data Availability

The infectious disease data were obtained from iedb.org. The cancer neoepitope data were obtained from cedar.iedb.org. The cancer survival data were obtained as previously described [37]. Data are freely available via Github at https://github.com/KrishnaswamyLab/ImmunoStruct.

## Code Availability

Source code for the ImmunoStruct model and inference scripts are available under an open-source license at https://github.com/KrishnaswamyLab/ImmunoStruct.

## Acknowledgements

This work was supported by the National Science Foundation (NSF Career Grant 2047856, NSF IIS 2473317, NSF DMS 2327211) and the National Institute of Health (NIH 1R01GM130847-01A1, NIH 1R01GM135929-01).

## Author Contributions

K.B.G., A.I. and S.K. identified the research problem and designed this work. K.B.G. collected and cleaned the IEDB and cancer data. K.B.G., J.F.R., E.Y., C.L., A.I. and S.K. conceived the experiments. K.B.G., J.F.R., E.Y. and C.L. developed ImmunoStruct. C.L. and K.B.G conceived, designed and developed the cancer-wildtype contrastive learning. K.B.G., J.F.R., E.Y. and C.L. trained and evaluated ImmunoStruct. K.B.G, K.G. and E.C. performed the experimental validation on SARS-CoV-2. K.B.G., J.F.R. and C.L. performed the data analysis. K.B.G. and C.L. produced the figures. All authors participated in the discussion and wrote the manuscript.

## Supplementary Materials

**Figure S1:**
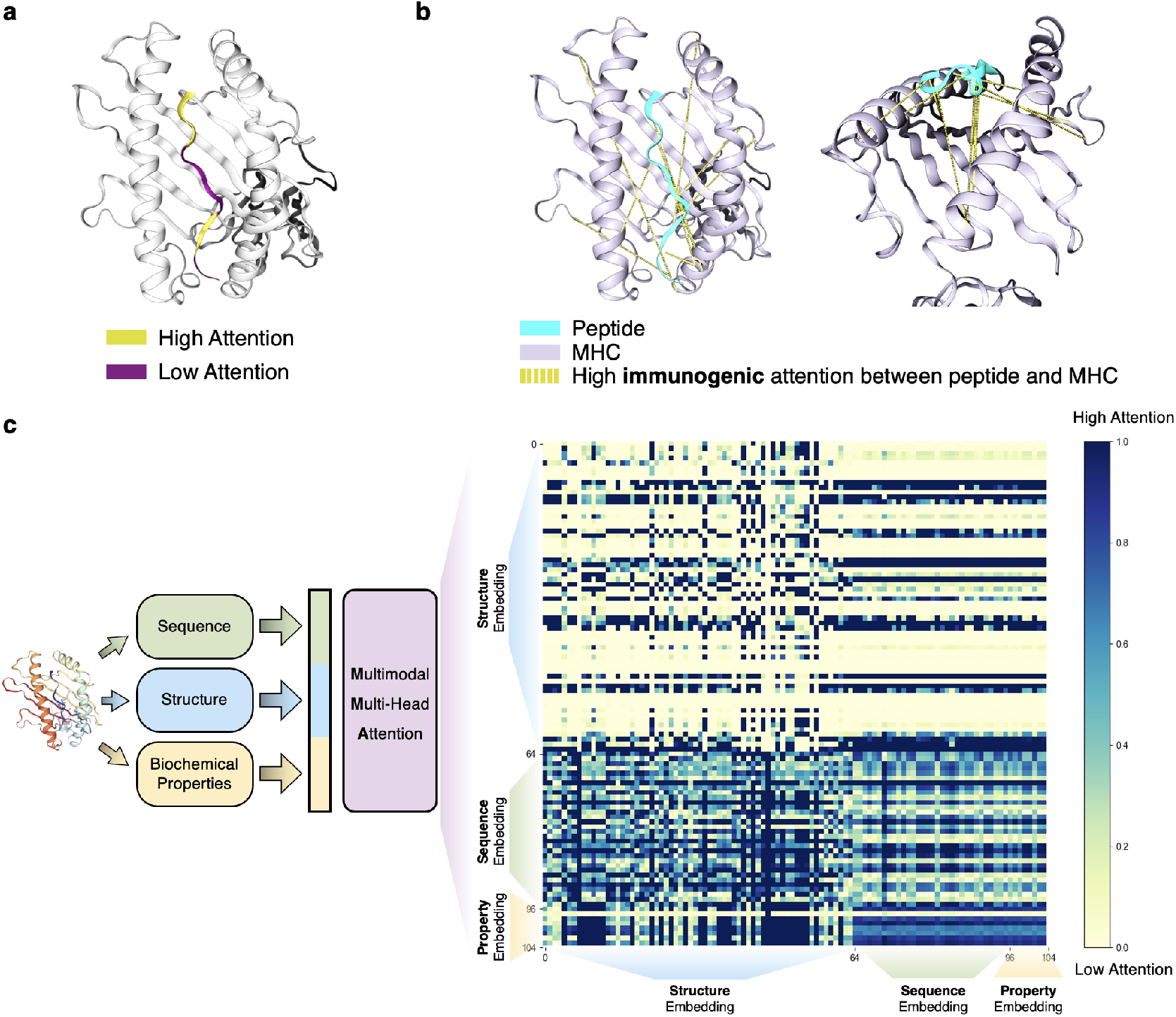
ImmunoStruct attention visualization examples on the peptide-MHC structure, and for the multi-modal embeddings. (a) The EGNN module attention layer can be extracted an visualized for both high (yellow) and low (purple) attention weights, directly on the peptide docked into the MHC protein. (b) The EGNN node-to-node attention weights between the peptide and the MHC protein can also be visualized as dotted lines (yellow) between the docked peptide and MHC protein. (c) The MMA learned attention matrix visualized as a heatmap across embeddings generated from the sequence, structure, and biochemical property data modalities.

**Figure S2:**
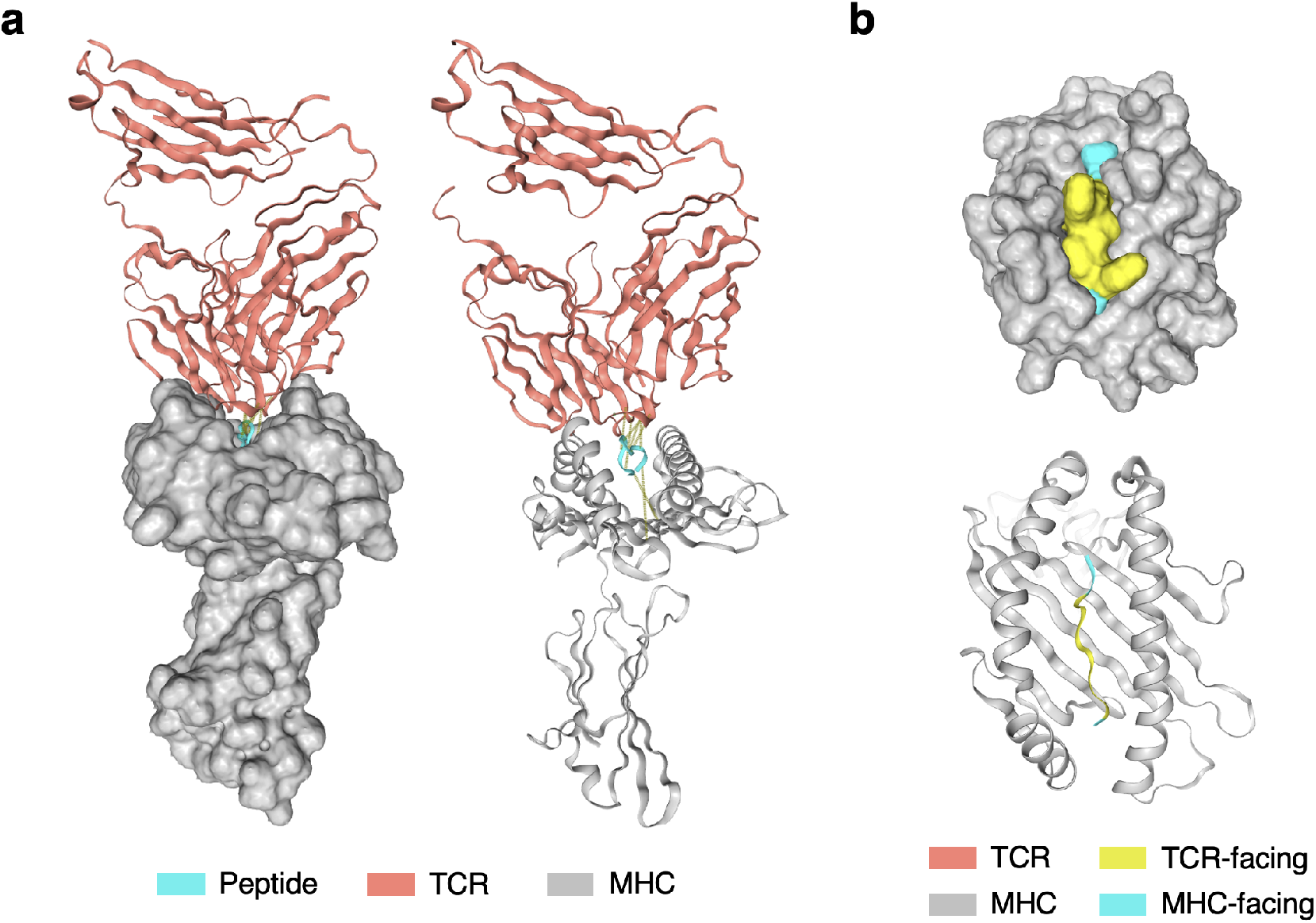
Additional structural illustrations of the MHC-facing and TCR-facing positions of the peptide. **(a)** Surface view (left) and cartoon view (right) of the TCR-peptide-MHC. **(b)** Zoom-in surface view (top) and cartoon view (bottom) of the peptide-MHC, with TCR-facing and MHC-facing segments of the peptide differently colored.

**Figure S3:**
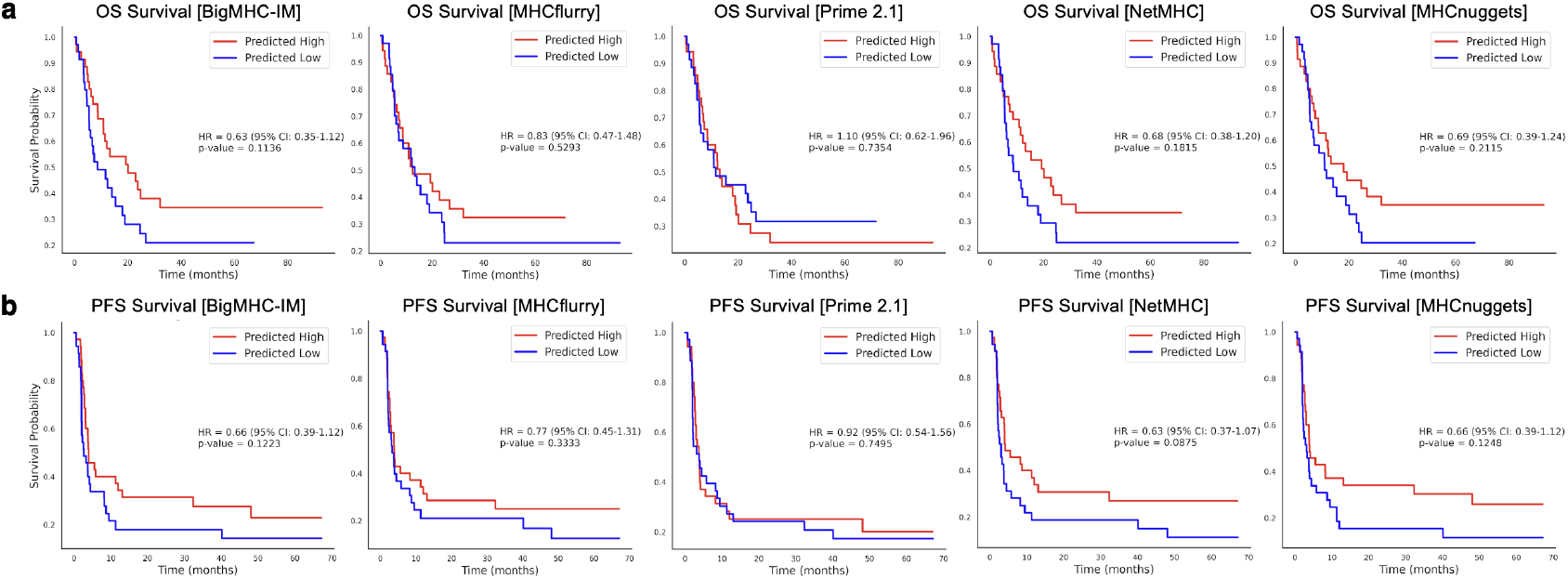
Kaplan-Meier survival plots for BigMHC, MHCflurry, Prime 2.1, NetMHC, and MHCnuggets showing these method’s respective stratification performance on the cancer clinical survival dataset for (a) Overall Survival (OS) and (b) Progression-free survival (PFS). The median split for predicted immunogenic peptide-MHC count was used.

**Table S1:** Ablation on the model components on **IEDB data**. All experiments were performed under 5 random seeds, and the mean and standard deviation (AUROC, AUPRC) or standard error of the mean (Mean PPVn) are displayed. For each metric, the best candidate is bolded. Bchems stands for biochemical properties and MMA stands for multimodal multi-head attention.

| Components |  |  |  | Transfer | AUROC | AUPRC | mean PPVn |
| --- | --- | --- | --- | --- | --- | --- | --- |
| Structure | Sequence | Bchems | MMA | Learning? |  |  |  |
| ✓ | ✓ | ✓ | ✓ | ✓ | **0.882** (0.005) | **0.696** (0.020) | **0.514** (0.021) |
| ✓ | ✓ | ✓ |  | ✓ | 0.840 (0.006) | 0.554 (0.028) | 0.441 (0.029) |
|  | ✓ | ✓ |  | ✓ | 0.844 (0.006) | 0.561 (0.017) | 0.442 (0.024) |
| ✓ | ✓ |  |  | ✓ | 0.844 (0.005) | 0.569 (0.014) | 0.440 (0.035) |
|  | ✓ |  |  | ✓ | 0.845 (0.007) | 0.568 (0.024) | 0.427 (0.017) |
| ✓ |  |  |  | ✓ | 0.805 (0.006) | 0.497 (0.022) | 0.414 (0.027) |
| ✓ | ✓ | ✓ | ✓ |  | 0.860 (0.013) | 0.615 (0.045) | 0.462 (0.015) |
| ✓ | ✓ | ✓ |  |  | 0.840 (0.006) | 0.547 (0.018) | 0.440 (0.032) |
|  | ✓ | ✓ |  |  | 0.842 (0.006) | 0.554 (0.020) | 0.419 (0.017) |
| ✓ | ✓ |  |  |  | 0.840 (0.007) | 0.547 (0.024) | 0.444 (0.028) |
|  | ✓ |  |  |  | 0.842 (0.006) | 0.553 (0.014) | 0.430 (0.009) |
| ✓ |  |  |  |  | 0.775 (0.012) | 0.442 (0.022) | 0.433 (0.020) |

**Table S2:** Comparison of immunogenicity prediction methods on **IEDB data**. All experiments were performed under 5 random seeds, and the mean and standard deviation (AUROC, AUPRC) or standard error of the mean (Mean PPVn) are displayed. For each metric, the best candidate is bolded.

| Model | AUROC | AUPRC | mean PPVn |
| --- | --- | --- | --- |
| Prime-2.1 [21] | **0.538** (0.012) | 0.212 (0.031) | 0.207 (0.013) |
| NetMHCpan [13] | **0.537** (0.027) | 0.214 (0.020) | 0.220 (0.008) |
| MHCnuggets [14] | **0.546** (0.023) | 0.246 (0.019) | 0.233 (0.014) |
| MHCflurry [15] | **0.577** (0.021) | 0.260 (0.026) | 0.273 (0.013) |
| DeepNeo [22] | **0.767** (0.032) | 0.438 (0.024) | 0.411 (0.008) |
| BigMHC-EL [16] | 0.588 (0.018) | 0.269 (0.016) | 0.290 (0.011) |
| BigMHC-IM [16] | 0.684 (0.028) | 0.319 (0.048) | 0.333 (0.015) |
| BigMHC<sub>retrained</sub> [16] | **0.793** (0.013) | 0.462 (0.027) | 0.445 (0.004) |
| **ImmunoStruct (ours)** | **0.882** (0.005) | **0.696** (0.020) | **0.514** (0.009) |

**Table S3:** Comparison of immunogenicity prediction methods on **cancer neoepitopes**. All experiments were performed under 5 random seeds, and the mean and standard deviation (AUROC, AUPRC) or standard error of the mean (Mean PPVn). For each metric, the best candidate is bolded.

| Model | AUROC | AUPRC | mean PPVn |
| --- | --- | --- | --- |
| Prime-2.1 [21] | 0.645 (0.026) | 0.259 (0.023) | 0.295 (0.013) |
| NetMHCpan [13] | 0.559 (0.049) | 0.210 (0.037) | 0.211 (0.020) |
| MHCnuggets [14] | 0.530 (0.014) | 0.210 (0.017) | 0.175 (0.010) |
| MHCflurry [15] | 0.658 (0.015) | 0.304 (0.032) | 0.329 (0.013) |
| DeepNeo [22] | 0.535 (0.016) | 0.261 (0.053) | 0.222 (0.031) |
| BigMHC-EL [16] | 0.632 (0.034) | 0.248 (0.039) | 0.245 (0.019) |
| BigMHC-IM [16] | **0.771** (0.040) | **0.373** (0.062) | **0.357** (0.016) |
| BigMHC<sub>retrained</sub> [16] | 0.682 (0.012) | 0.310 (0.020) | 0.325 (0.013) |
| **ImmunoStruct (ours)** | **0.771** (0.024) | **0.433** (0.069) | **0.364** (0.057) |

## References

[1] Madhu Gupta, Abhishek Wahi, Priyanka Sharma, Riya Nagpal, Neha Raina, Monika Kaurav, Jaydeep Bhattacharya, Sonia M Rodrigues Oliveira, Karma G Dolma, Alok K Paul, et al. Recent advances in cancer vaccines: challenges, achievements, and futuristic prospects. Vaccines, 10(12):2011, 2022.

[2] Ting Fan, Mingna Zhang, Jingxian Yang, Zhounan Zhu, Wanlu Cao, and Chunyan Dong. Therapeutic cancer vaccines: advancements, challenges, and prospects. Signal Transduction and Targeted Therapy, 8(1):450, 2023.

[3] Eryn Blass and Patrick A Ott. Advances in the development of personalized neoantigen-based therapeutic cancer vaccines. Nature reviews Clinical oncology, 18(4):215–229, 2021.

[4] Anne-Mette Bjerregaard, Morten Nielsen, Vanessa Jurtz, Carolina M Barra, Sine Reker Hadrup, Zoltan Szallasi, and Aron Charles Eklund. An analysis of natural t cell responses to predicted tumor neoepitopes. Frontiers in immunology, 8:1566, 2017.

[5] Peter D Katsikis, Ken J Ishii, and Christopher Schliehe. Challenges in developing personalized neoantigen cancer vaccines. Nature Reviews Immunology, 24(3):213–227, 2024.

[6] Patrick A Ott, Siwen Hu-Lieskovan, Bartosz Chmielowski, Ramaswamy Govindan, Aung Naing, Nina Bhardwaj, Kim Margolin, Mark M Awad, Matthew D Hellmann, Jessica J Lin, et al. A phase ib trial of personalized neoantigen therapy plus anti-pd-1 in patients with advanced melanoma, non-small cell lung cancer, or bladder cancer. Cell, 183(2):347–362, 2020.

[7] Daniel K Wells, Marit M van Buuren, Kristen K Dang, Vanessa M Hubbard-Lucey, Kathleen CF Sheehan, Katie M Campbell, Andrew Lamb, Jeffrey P Ward, John Sidney, Ana B Blazquez, et al. Key parameters of tumor epitope immunogenicity revealed through a consortium approach improve neoantigen prediction. Cell, 183(3):818–834, 2020.

[8] Ugur Sahin, Evelyna Derhovanessian, Matthias Miller, Björn-Philipp Kloke, Petra Simon, Martin Löwer, Valesca Bukur, Arbel D Tadmor, Ulrich Luxemburger, Barbara Schrörs, et al. Personalized rna mutanome vaccines mobilize poly-specific therapeutic immunity against cancer. Nature, 547(7662): 222–226, 2017.

[9] Novalia Pishesha, Thibault J Harmand, and Hidde L Ploegh. A guide to antigen processing and presentation. Nature Reviews Immunology, 22(12):751–764, 2022.

[10] Fleur E Tynan, Diah Elhassen, Anthony W Purcell, Jacqueline M Burrows, Natalie A Borg, John J Miles, Nicholas A Williamson, Kate J Green, Judy Tellam, Lars Kjer-Nielsen, et al. The immunogenicity of a viral cytotoxic t cell epitope is controlled by its mhc-bound conformation. The Journal of experimental medicine, 202(9):1249–1260, 2005.

[11] Peng Wu, Tongtong Zhang, Baoyu Liu, Panyu Fei, Lei Cui, Rui Qin, Huaying Zhu, Danmei Yao, Ryan J Martinez, Wei Hu, et al. Mechano-regulation of peptide-mhc class i conformations determines tcr antigen recognition. Molecular cell, 73(5):1015–1027, 2019.

[12] Jeffrey K Weber, Joseph A Morrone, Seung-gu Kang, Leili Zhang, Lijun Lang, Diego Chowell, Chirag Krishna, Tien Huynh, Prerana Parthasarathy, Binquan Luan, et al. Unsupervised and supervised ai on molecular dynamics simulations reveals complex characteristics of hla-a2-peptide immunogenicity. Briefings in Bioinformatics, 25(1):bbad504, 2024.

[13] Birkir Reynisson, Bruno Alvarez, Sinu Paul, Bjoern Peters, and Morten Nielsen. Netmhcpan-4.1 and netmhciipan-4.0: improved predictions of mhc antigen presentation by concurrent motif deconvolution and integration of ms mhc eluted ligand data. Nucleic acids research, 48(W1):W449–W454, 2020.

[14] Xiaoshan M Shao, Rohit Bhattacharya, Justin Huang, IK Ashok Sivakumar, Collin Tokheim, Lily Zheng, Dylan Hirsch, Benjamin Kaminow, Ashton Omdahl, Maria Bonsack, et al. High-throughput prediction of mhc class i and ii neoantigens with mhcnuggets. Cancer immunology research, 8(3):396–408, 2020.

[15] Timothy J O’Donnell, Alex Rubinsteyn, and Uri Laserson. Mhcflurry 2.0: improved pan-allele prediction of mhc class i-presented peptides by incorporating antigen processing. Cell systems, 11(1):42–48, 2020.

[16] Benjamin Alexander Albert, Yunxiao Yang, Xiaoshan M Shao, Dipika Singh, Kellie N Smith, Valsamo Anagnostou, and Rachel Karchin. Deep neural networks predict class i major histocompatibility complex epitope presentation and transfer learn neoepitope immunogenicity. Nature machine intelligence, 5(8): 861–872, 2023.

[17] Kei Saotome, Drew Dudgeon, Kiersten Colotti, Michael J Moore, Jennifer Jones, Yi Zhou, Ashique Rafique, George D Yancopoulos, Andrew J Murphy, John C Lin, et al. Structural analysis of cancer-relevant tcr-cd3 and peptide-mhc complexes by cryoem. Nature Communications, 14(1):2401, 2023.

[18] Randi Vita, Shuchi Mahajan, James A. Overton, Sandeep K. Dhanda, Stefania Martini, Jeffrey R. Cantrell, Donna K. Wheeler, Alessandro Sette, and Bjoern Peters. The immune epitope database (iedb): 2018 update. Nucleic Acids Research, October 24 2018. doi: 10.1093/nar/gky1006.

[19] Zeynep Koşaloğlu-Yalçin, Nina Blazeska, Randi Vita, Hannah Carter, Morten Nielsen, Stephen Schoenberger, Alessandro Sette, and Bjoern Peters. The cancer epitope database and analysis resource (cedar). Nucleic acids research, 51(D1):D845–D852, 2023.

[20] John Jumper, Richard Evans, Alexander Pritzel, Tim Green, Michael Figurnov, Olaf Ronneberger, Kathryn Tunyasuvunakool, Russ Bates, Augustin Žídek, Anna Potapenko, et al. Highly accurate protein structure prediction with alphafold. nature, 596(7873):583–589, 2021.

[21] David Gfeller, Julien Schmidt, Giancarlo Croce, Philippe Guillaume, Sara Bobisse, Raphael Genolet, Lise Queiroz, Julien Cesbron, Julien Racle, and Alexandre Harari. Improved predictions of antigen presentation and tcr recognition with mixmhcpred2. 2 and prime2. 0 reveal potent sars-cov-2 cd8+ t-cell epitopes. Cell Systems, 14(1):72–83, 2023.

[22] Jeong Yeon Kim, Hyoeun Bang, Seung-Jae Noh, and Jung Kyoon Choi. Deepneo: a webserver for predicting immunogenic neoantigens. Nucleic acids research, 51(W1):W134–W140, 2023.

[23] Victor Garcia Satorras, Emiel Hoogeboom, and Max Welling. E (n) equivariant graph neural networks. In International conference on machine learning, pages 9323–9332. PMLR, 2021.

[24] Thomas N. Kipf and Max Welling. Semi-Supervised Classification with Graph Convolutional Networks. In International Conference on Learning Representations, 2017.

[25] Ashish Vaswani, Noam Shazeer, Niki Parmar, Jakob Uszkoreit, Llion Jones, Aidan N Gomez, Lukasz Kaiser, and Illia Polosukhin. Attention is all you need. Advances in Neural Information Processing Systems, 2017.

[26] Diederik P Kingma. Auto-encoding variational bayes. In International Conference on Learning Representations, 2014.

[27] Chen Liu, Danqi Liao, Alejandro Parada-Mayorga, Alejandro Ribeiro, Marcello DiStasio, and Smita Krishnaswamy. Diffkillr: Killing and recreating diffeomorphisms for cell annotation in dense microscopy images. In IEEE International Conference on Acoustics, Speech and Signal Processing (ICASSP). IEEE, 2025.

[28] Xingzhi Sun, Danqi Liao, Kincaid MacDonald, Yanlei Zhang, Chen Liu, Guillaume Huguet, Guy Wolf, Ian Adelstein, Tim GJ Rudner, and Smita Krishnaswamy. Geometry-aware generative autoencoders for warped riemannian metric learning and generative modeling on data manifolds. In International Conference on Artificial Intelligence and Statistics. PMLR, 2024.

[29] Daniel Osorio, Paola Rondón-Villarreal, and Rodrigo Torres. Peptides: a package for data mining of antimicrobial peptides. The R Journal, 7(1):4–14, 2015. doi: 10.32614/RJ-2015-001.

[30] Fuzhen Zhuang, Zhiyuan Qi, Keyu Duan, Dongbo Xi, Yongchun Zhu, Hengshu Zhu, Hui Xiong, and Qing He. A comprehensive survey on transfer learning. Proceedings of the IEEE, 109(1):43–76, 2020.

[31] Lee P. Richman, Robert H. Vonderheide, and Andrew J. Rech. Neoantigen dissimilarity to the self-proteome predicts immunogenicity and response to immune checkpoint blockade. Cell Systems, 9(4): 375–382, 2019.

[32] Marta Luksza, Nadeem Riaz, Vladimir Makarov, Vinod P. Balachandran, Matthew D. Hellmann, Alexander Solovyov, Naiyer A. Rizvi, Taha Merghoub, Arnold J. Levine, Timothy A. Chan, Jedd D. Wolchok, and Benjamin D. Greenbaum. A neoantigen fitness model predicts tumour response to checkpoint blockade immunotherapy. Nature, 551(7681):517–520, November 2017. ISSN 1476-4687. doi: 10.1038/nature24473. URL http://dx.doi.org/10.1038/nature24473.

[33] Vinod P Balachandran, Marta Luksza, Julia N Zhao, Vladimir Makarov, John Alec Moral, Romain Remark, Brian Herbst, Gokce Askan, Umesh Bhanot, Yasin Senbabaoglu, et al. Identification of unique neoantigen qualities in long-term survivors of pancreatic cancer. Nature, 551(7681):512–516, 2017.

[34] Justin SA Perry and Chyi-Song Hsieh. Development of t-cell tolerance utilizes both cell-autonomous and cooperative presentation of self-antigen. Immunological reviews, 271(1):141–155, 2016.

[35] Kevin R Moon, David Van Dijk, Zheng Wang, Scott Gigante, Daniel B Burkhardt, William S Chen, Kristina Yim, Antonia van den Elzen, Matthew J Hirn, Ronald R Coifman, et al. Visualizing structure and transitions in high-dimensional biological data. Nature biotechnology, 37(12):1482–1492, 2019.

[36] Daniel M Tadros, Simon Eggenschwiler, Julien Racle, and David Gfeller. The mhc motif atlas: a database of mhc binding specificities and ligands. Nucleic acids research, 51(D1):D428–D437, 2023.

[37] Annie Borch, Ibel Carri, Birkir Reynisson, Heli M Garcia Alvarez, Kamilla K Munk, Alessandro Montemurro, Nikolaj Pagh Kristensen, Siri A Tvingsholm, Jeppe Sejerø Holm, Christina Heeke, et al. Improve: a feature model to predict neoepitope immunogenicity through broad-scale validation of t-cell recognition. Frontiers in Immunology, 15:1360281, 2024.

[38] Amy R Rappaport, Chrisann Kyi, Monica Lane, Meghan G Hart, Melissa L Johnson, Brian S Henick, Chih-Yi Liao, Amit Mahipal, Ardaman Shergill, Alexander I Spira, et al. A shared neoantigen vaccine combined with immune checkpoint blockade for advanced metastatic solid tumors: phase 1 trial interim results. Nature Medicine, 30(4):1013–1022, 2024.

[39] Ji-Li Chen, Guillaume Stewart-Jones, Giovanna Bossi, Nikolai M Lissin, Linda Wooldridge, Ed Man Lik Choi, Gerhard Held, P Rod Dunbar, Robert M Esnouf, Malkit Sami, et al. Structural and kinetic basis for heightened immunogenicity of t cell vaccines. The Journal of experimental medicine, 201(8):1243–1255, 2005.

[40] Daniela Weiskopf, Michael A Angelo, Elzinandes L de Azeredo, John Sidney, Jason A Greenbaum, Anira N Fernando, Anne Broadwater, Ravi V Kolla, Aruna D De Silva, Aravinda M de Silva, et al. Comprehensive analysis of dengue virus-specific responses supports an hla-linked protective role for cd8+ t cells. Proceedings of the National Academy of Sciences, 110(22):E2046–E2053, 2013.

[41] Marta Luksza, Nadeem Riaz, Vladimir Makarov, Vinod P Balachandran, Matthew D Hellmann, Alexander Solovyov, Naiyer A Rizvi, Taha Merghoub, Arnold J Levine, Timothy A Chan, et al. A neoantigen fitness model predicts tumour response to checkpoint blockade immunotherapy. Nature, 551 (7681):517–520, 2017.

[42] Geoffrey E Hinton and Ruslan R Salakhutdinov. Reducing the dimensionality of data with neural networks. science, 313(5786):504–507, 2006.

[43] Chen Liu, Ke Xu, Liangbo L Shen, Guillaume Huguet, Zilong Wang, Alexander Tong, Danilo Bzdok, Jay Stewart, Jay C Wang, Lucian V Del Priore, and Smita Krishnaswamy. Imageflownet: Forecasting multiscale trajectories of disease progression with irregularly-sampled longitudinal medical images. In IEEE International Conference on Acoustics, Speech and Signal Processing (ICASSP). IEEE, 2025.

[44] Chen Liu, Matthew Amodio, Liangbo L Shen, Feng Gao, Arman Avesta, Sanjay Aneja, Jay C Wang, Lucian V Del Priore, and Smita Krishnaswamy. Cuts: A deep learning and topological framework for multigranular unsupervised medical image segmentation. In International Conference on Medical Image Computing and Computer-Assisted Intervention, pages 155–165. Springer, 2024.

[45] Adam Paszke, Sam Gross, Francisco Massa, Adam Lerer, James Bradbury, Gregory Chanan, Trevor Killeen, Zeming Lin, Natalia Gimelshein, Luca Antiga, et al. Pytorch: An imperative style, high-performance deep learning library. Advances in neural information processing systems, 32, 2019.

[46] Ting Chen, Simon Kornblith, Mohammad Norouzi, and Geoffrey Hinton. A simple framework for contrastive learning of visual representations. In International conference on machine learning, pages 1597–1607. PMLR, 2020.

[47] Jure Zbontar, Li Jing, Ishan Misra, Yann LeCun, and Stéphane Deny. Barlow twins: Self-supervised learning via redundancy reduction. In International conference on machine learning, pages 12310–12320. PMLR, 2021.

[48] Ilya Loshchilov and Frank Hutter. Decoupled weight decay regularization. In International Conference on Learning Representations, 2019.

[49] Ilya Loshchilov and Frank Hutter. Sgdr: Stochastic gradient descent with warm restarts. arXiv preprint 1608.03983, 2016.

[50] Alessandro Sette and John Sidney. Nine major hla class i supertypes account for the vast preponderance of hla-a and -b polymorphism. Immunogenetics, 50(3):201–212, 1999.

[51] Milot Mirdita, Konstantin Schütze, Yoshitaka Moriwaki, Lim Heo, Sergey Ovchinnikov, and Martin Steinegger. Colabfold: making protein folding accessible to all. Nature methods, 19(6):679–682, 2022.

[52] Neeha Zaidi, Mariya Soban, Fangluo Chen, Heather Kinkead, Jocelyn Mathew, Mark Yarchoan, Todd D Armstrong, Shozeb Haider, and Elizabeth M Jaffee. Role of in silico structural modeling in predicting immunogenic neoepitopes for cancer vaccine development. JCI insight, 5(17), 2020.

[53] Meredith Slota, Jong-Baeck Lim, Yushe Dang, and Mary L. Disis. Elispot for measuring human immune responses to vaccines. Expert Review of Vaccines, 10(3):299–306, 2011. doi: 10.1586/erv.10.169.

[54] Fan Yang, Kathryn Patton, Theresa Kasprzyk, Brain Long, Soumi Gupta, Stephen J. Zoog, Kristin Tracy, and Christian Vettermann. Validation of an ifn-gamma elispot assay to measure cellular immune responses against viral antigens in non-human primates. Gene Therapy, 29(1-2):41–54, 2022. doi: 10.1038/s41434-020-00214-w.

[55] A. Nelde, T. Bilich, J.S. Heitmann, et al. Sars-cov-2-derived peptides define heterologous and covid-19-induced t cell recognition. Nature Immunology, 22(1):74–85, 2021. doi: 10.1038/s41590-020-00808-x.

